# Transcranial direct current stimulation combined cognitive training modulates risk-taking behavior in older adults

**DOI:** 10.64898/2026.02.28.708700

**Authors:** Ping Ren, Yunchen Gong, Manxiu Ma, Yu Fu, Yuchuan Zhuang, Donghui Wu, Li Zhang

## Abstract

Risk-taking behavior is a complex cognitive function that often declines with aging, contributing to impaired decision-making and reduced quality of life. While transcranial direct current stimulation (tDCS) has shown promising effects in modulating cognitive function, its influence on ecologically valid, complex behaviors like risky decision-making in older adults remains poorly understood. We investigated whether a single-session stimulation over the medial orbitofrontal cortex (MOFC) combined with cognitive training could enhance decision-making in healthy older adults. In a randomized, sham-controlled study, bilateral MOFC stimulation (left/anode, right/cathode) was applied during a training task based on the Iowa Gambling Task. Pre- and post-intervention assessments utilized conventional behavioral measures and the Values-Plus-Perseveration computational model, alongside task-related fMRI to examine MOFC network changes. Compared to sham, tDCS significantly enhanced the ability to distinguish advantageous from disadvantageous options. Modeling analysis revealed stimulation-induced changes in multiple latent components, such as learning rate, loss aversion, and perseveration decay. Generalized Psychophysiological Interaction analysis showed that tDCS reconfigured the MOFC network by reducing fronto-frontal hyper-connectivity and enhancing fronto-striatal connectivity. These behavioral improvements were specifically associated with the left MOFC network targeted by anodal stimulation. Our results provide causal evidence that tDCS can mitigate age-related impairments in risk-taking by reconfiguring MOFC-related networks. These findings advance the mechanistic understanding of tDCS in aging and highlight its potential as an intervention to prevent age-related decline in decision making.

## 1. Introduction

Decision making under uncertainty is critical for individual’s survival and development, which is tightly intertwined with risk. To cope in uncertain environments, individuals need to optimize their decision strategies by maximizing gains and minimizing losses. Impaired risky decision-making has been found in many neuropsychiatric disorders, as well as in normal and pathologic aging (Dalley and Robbins, 2017; Frank and Seaman, 2023; Samanez-Larkin and Knutson, 2015).

Compared with young adults, elderly people often have difficulty in making decisions under uncertainty, increasing the risk of financial exploitation. Although neuroplasticity declines substantially with aging, an increasing number of studies have reported promising effects of non-pharmacological interventions, such as non-invasive brain stimulation (NIBS) and cognitive training, in improving cognitive performance in older adults (Aksu et al., 2024; Antonenko et al., 2024; Indahlastari et al., 2020; Tang et al., 2019). Nevertheless, studies applying tDCS to complex behaviors such as risky decision-making remain scarce, especially in the elderly population.

The Iowa Gambling Task (IGT), in which probabilities and possible outcomes need to be learned through feedback from previous choices, has been widely used to investigate individual’s decision process under risk (Hultman et al., 2022; Zanini et al., 2025). The test-retest reliability and validity of the IGT have been confirmed in assessing decision making in both healthy and clinical populations (Cardoso et al., 2010; Hasuzawa et al., 2022; Pasion et al., 2017; Xu et al., 2013). In aging-related studies, IGT net score, defined by advantageous versus disadvantageous choices, has been found significantly reduced in older adults compared with young adults (Di Rosa et al., 2017; Fein et al., 2007; Ren et al., 2024). In addition to conventional behavioral metrics, a series of IGT computational models have been developed to describe decision making process, such as the Expectancy–Valence Learning model (the first proposed model for the IGT) (Busemeyer and Stout, 2002), the Prospect Valence Learning model with Delta (PVL-Delta) (Ahn et al., 2008), the PVL model with decay (PVL-Decay) (Ahn et al., 2014), and the Values-Plus-Perseveration (VPP) model (Worthy et al., 2013). Compared with other three models, the VPP model provides eight parameters, showing more informative components during the IGT by decomposing perseverative tendency and expected value representation. A number of recent studies have successfully applied the VPP model to assess alterations of IGT performance in chronic pain (Zhang et al., 2022), methamphetamine use disorder (Liu et al., 2024), autism spectrum disorder and obsessive-compulsive disorder (Carlisi et al., 2017). Therefore, integrating conventional behavioral measures with computational modeling parameters would offer deeper insights into age-related changes in decision-making processes.

The medial orbitofrontal cortex (MOFC), which is anatomically connected to the striatum (Park et al., 2022), has been found closely associated with value-based decision-making (Lopez-Persem et al., 2020). Combining functional magnetic resonance imaging (fMRI) and machine-learning tools, multiple voxels activity pattern in the orbitofrontal areas, especially the MOFC, represent advantageous choice versus disadvantageous choice in the IGT (Zha et al., 2022). Recently, our previous study has reported that disruption of MOFC-centered circuitry may lead to altered risk-taking behavior in older adults (Ren et al., 2024). Therefore, we speculated that the MOFC network plays a pivotal role in alterations of risky decision-making in aging and might be a potential target for NIBS-based intervention.

Transcranial direct current stimulation (tDCS), one of the most popular NIBS techniques, has been widely used in fundamental research and clinical treatment (Dedoncker et al., 2016; Summers et al., 2016). The primary advantages of tDCS are the absence of serious side effects and the portability of the devices, making it especially appropriate for older adults. Numerous studies have demonstrated that tDCS can effectively modulate cortical activity using weak electrical currents; specifically, anodal stimulation increases, whereas cathodal stimulation decreases, cortical excitability (Mordillo-Mateos et al., 2012; Pellicciari et al., 2013; Zheng et al., 2011). Notably, prefrontal tDCS has been found to enhance neural activity in subcortical components of the dopaminergic system, including the striatum, suggesting that its effects extend beyond the stimulation site. (Meyer et al., 2019; Weber et al., 2014). In healthy young adults, a behavioral study has demonstrated that tDCS applied bilaterally over the orbitofrontal cortex could improve IGT net score (Ouellet et al., 2015). However, it remains under-investigated whether and how tDCS over the MOFC might improve risk-taking behavior in older adults.

In the present study, we investigated the MOFC-tDCS combining cognitive training task in modulating risky decision-making and underlying brain networks in healthy elderly people. Given the complexity and variation of cognitive deterioration in aging, we analyzed both conventional task measures and VPP model components to better characterize IGT performance before and after intervention. We hypothesized that tDCS over the bilateral MOFC would effectively change IGT net score and model parameters in elderlies. Combining task-related fMRI analysis, we expected that the MOFC would interplay with subcortical regions such as striatum to regulate risk-taking behavior in response to current stimulation. Specifically, the aims of the present study are three-folds: (1) to examine the MOFC-tDCS effect on risk-taking behavior in older adults; (3) to determine whether and how the MOFC network changes in response to tDCS intervention. (2) to validate the relationship between intervention-induced changes in MOFC connectivity and changes in risk-taking behavior.

## 2. Material and Methods

### 2.1 Participants

Fifty-two healthy older adults (aged 56 – 88) were recruited from multiple communities in Shenzhen. All participants were right-handed determined by the Edinburgh handedness inventory, have adequate visual and auditory acuity for testing by self-report. The individuals with one of the following conditions were excluded: (1) a history of diagnosed neurological or psychiatric diseases such as cerebrovascular disease and major depression; (2) MRI contraindications (i.e., metallic implant, claustrophobia, or pacemaker); (3) the Mini-mental State Examination (MMSE) score ≤ 20. All participants were required to sign a written informed consent form after a full written and verbal explanation of the study. This study was performed in accordance with the Declaration of Helsinki and had been approved by the Ethics Committee of Shenzhen Mental Health Center/Shenzhen Kangning Hospital.

### 2.2 Neuropsychological measurement

For each participant, global cognitive function was assessed using the 30-point MMSE, a commonly used screening tool to assess multiple cognitive domains, including orientation, short-term memory, spatial abilities and so on (Folstein et al., 1975). The Hamilton Rating Scale for Depression (HAMD) was used to assess the severity of Depression, and individuals with HAMD > 16 (moderate to severe symptom) were excluded from the analysis (Zimmerman et al., 2013). To minimize the influence of cognitive and emotional states on risky decision-making, the MMSE and HAMD scores were controlled as covariates in the following analysis.

### 2.3 Iowa Gambling Task

A modified IGT, based on Cauffman and colleague’s version (Cauffman et al., 2010), was delivered to assess risk-taking behavior during scanning. This version has been successfully used to examine age-related alterations of risky decision-making in our previous studies (Ren et al., 2023; Ren et al., 2024). Briefly, each trial started with a yellow frame on one of the four decks randomly, and participants need to choose ‘play’ or ‘pass’ the card by pressing button in their left and right hands (**Figure 1A**). Out of every ten cards, deck A gave 50% chance of losing ¥95, deck B gave 20% chance of losing ¥115, deck C gave 50% chance of losing ¥35, and deck D gave 20% chance of losing ¥25. Therefore, advantageous choices (deck C and D) would give participants a cumulative gain, while disadvantageous choices (deck A and B) would give a loss. During the task, participants need to learn the covert rule and maximize their gains. Then a net score, calculated by subtracting disadvantageous choices from advantageous choices, was used to assess participant’s decision-making ability.

**Figure 1.**
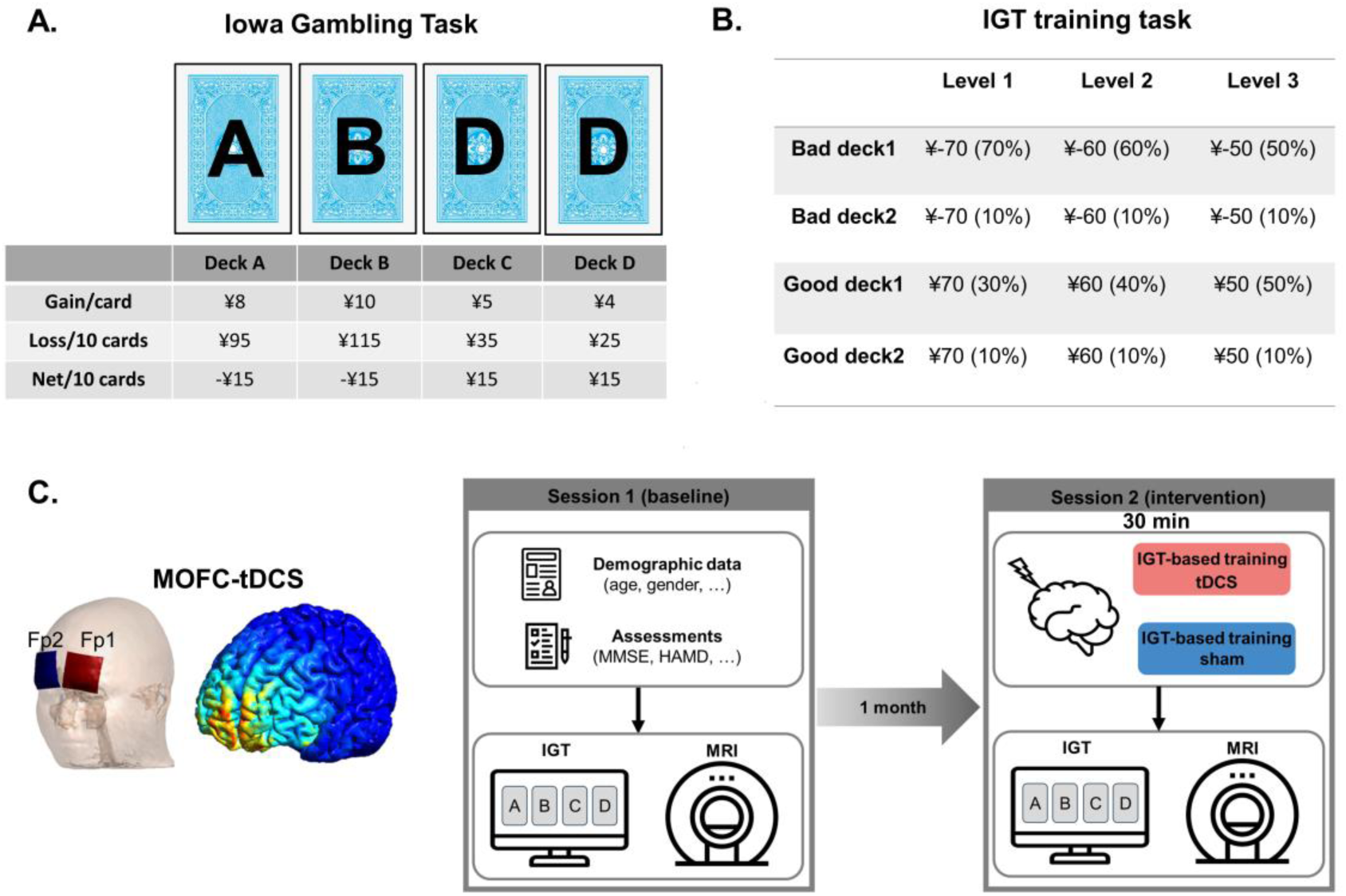
Iowa gambling task and tDCS experimental procedure. A) The IGT and net gain/loss parameters. B) The IGT-based cognitive training paradigm was consist of three blocks with different difficulty levels. C) In the tDCS, anodal/cathodal electrodes were placed over the left and right MOFC (Fp1/Fp2). Simulated electric field induced by tDCS was computed with SimNIBS. In the interventional session, participant was required to complete the tDCS combining IGT training task and a MRI scanning.

### 2.4 Value-plus-preservation model

In addition to conventional behavioral analysis, the VPP model provides a more informative components of the IGT performance by estimating latent decision making process (Worthy et al., 2013). Based on the Bayesian logic, the VPP model delineates eight parameters: memory decay (*A*) represents the discount for past expected values. A higher value indicates stronger influence of past outcomes in the valuation process. Learning rate (*α*) controls how much an individual updates their belief about an option’s value after receiving feedback. A higher value indicates greater sensitivity to objective value changes. Consistency (*cons*) signifies response consistency. Higher consistency means more deterministic choice, while lower consistency means more exploratory behavior. Loss aversion (*λ*) denotes sensitivity to gain and loss. *λ* > 1 indicates that the individual’s behavior is more sensitive to losses than to gains. Positive and negative outcome preservation (*epP* and *epN*) measures preservation strength in response to gain/loss. Larger *epP* indicates stronger repetition after gains, while positive *epN* indicates loss-stay behavior and negative *epN* indicates loss-shift behavior. Perseveration decay (*K*) decides the decay rate of the persistence strength of all decks on each trial. A higher value means a stronger influence of the previous perseverance tendency. Reinforcement learning rate (*w*) balances expected valence and perseverance tendency on decisions. A higher value of *w* indicates a stronger influence of expected valence.

### 2.5 Cognitive training task

In the second session, each participant completed a cognitive training task during tDCS. Building on the conventional IGT, we developed a training paradigm consisting of three blocks with increasing difficulty levels (**Figure 3B**). In the task, participants chose between two bad decks (risky options) and two good decks (safe options), as defined in the conventional IGT. In Level 1, Bad Deck 1 produced a loss of ¥70 on 70% of trials, while Bad Deck 2 produced a loss of ¥70 on 10% of trials. Good Deck 1 yielded a reward of ¥70 on 30% of trials, and Good Deck 2 yielded a reward of ¥70 on 10% of trials. In Level 2, the differences in reward/loss probabilities between the advantageous and disadvantageous decks were reduced, requiring greater effort to optimize strategy. In Level 3, these differences were reduced further, increasing task difficulty. The entire training lasted 30 minutes, divided into three consecutive 10-minute blocks. This progressively challenging paradigm helped older participants detect and learn the underlying rules, thereby facilitating strategy optimization throughout the task.

### 2.6 Experimental procedure

Current stimulation was delivered with a battery-driven DC stimulator (DROIAN2019) with two saline-soaked surface sponge electrodes (current 1.5 mA, area 5×5 cm^2^, intensity 60 μA/cm^2^). According to the EEG 10-20 international system, the anodal electrode was placed on the left MOFC (Fp1), and the cathodal electrode was placed on the right MOFC (Fp2). In the tDCS sessions, the current was administered for 30 min, with a 20 second ramp in and ramp out. In the sham session, the current was administered for 1 min with a 20 second ramp in and ramp out, to ensure that participants experienced a similar initial sensation as in real stimulation. All participants were kept blinded to the stimulation conditions throughout study.

All participants were required to complete two experimental sessions, without taking alcohol or caffeinated drink before each session. In Session 1 (baseline screening), demographic data and neuropsychological assessments were collected from each participant. Then participant was required to complete the IGT during MRI scanning. To minimize learning effect, Session 2 was conducted one-month after Session 1. In Session 2 (tDCS intervention), each participant was required to complete a 30-min cognitive training task with tDCS intervention (real or sham) using a single-blind design. Within 30 minutes following the tDCS intervention, participants underwent MRI scanning and completed the IGT. (**Figure 1D**).

### 2.7 Imaging data acquisition and preprocessing

In Session 1 and 2, neuroimaging data were acquired on a 3.0T MRI system (Discovery MR750 System, GE Healthcare) with an eight-channel phased-array head coil. The T1-weighted structural images were acquired by using a three-dimensional brain volume imaging sequence that covered the whole brain (repetition time (TR) = 6.7 ms, echo time (TE) = 2.9 ms, flip angle = 7 degrees, matrix = 256×256, slice thickness = 1 mm, 196 slices). Then a task-related fMRI was delivered using gradient-echo echo-planar imaging sequence with the following parameters: TR = 2000 ms, TE = 25 ms, flip angle = 90 degrees, matrix = 64×64, and voxel size = 3.4×3.4×3.2 mm^3^, and 48 axial slices. During the task session, the IGT task was projected on the screen, and participants made responses by pressing left-handed button for play and right-handed button for pass. The entire task lasted 10 minutes with 300 volumes.

The functional images were preprocessed in the DPABI_v4.3 (https://rfmri.org/dpabi) (Yan et al., 2016) based on the SPM8 (http://www.fil.ion.ucl.ac.uk/spm/). For functional imaging data, the first 5 volumes were excluded to obtain steady-state tissue magnetization. The remaining volumes were corrected for slice timing and head motion, co-registered to their own structural images, and normalized to the Montreal Neurological Institute (MNI) standard space. Then the functional images were resampled to 3 × 3 × 3 mm, and smoothed with a Gaussian kernel (FWHM = 6 mm).

After data preprocessing step, head motion was carefully checked, due to the commonality of head motion in older adults and its confounding effect on resting-state functional connectivity (Power et al., 2012; Van Dijk et al., 2012). First, no participant exhibited head motion exceeding 2.5 mm in translation or 2.5 degrees in rotation during the task-related scanning. Second, framewise displacement (FD), a measure of head movement between consecutive volumes, was calculated as the sum of the absolute values of the derivatives of the six realignment parameters (Power et al., 2012). For each participant, mean FD did not exceed 0.5 mm in either session. In addition, FD-pre (Session 1) and FD-post (Session 2) were controlled as covariates of non-interest in functional connectivity analyses.

### 2.8 Gray matter atrophy

The structural images were preprocessed to generate a whole-brain gray matter map using Voxel-based morphometry (VBM). For each participant, the T1-weighted image was segmented into gray matter, white matter, and cerebrospinal fluid. Then, a gray matter template was generated through an iterative nonlinear registration using DARTEL, a toolbox with a fast diffeomorphic registration algorithm (Ashburner, 2007). For each participant, an averaged gray matter (GM) volume of the entire brain was computed to control age-induced brain atrophy in the MRI data analyses.

### 2.9 Generalized Psychophysiological Interaction (gPPI) analysis

A general linear model was applied to estimate BOLD signals corresponding to the four event-related regressors, representing deck A, B, C, and D. Regressors were generated by convolving the onset times of each event with a canonical hemodynamic response function. Non-interested covariates, including six head-motion parameters, white matter and CSF signals were included in the design matrix. Then a high-pass filter with a cut-off period of 128 seconds was used to remove slow drift. For each participant, trials without response were excluded, and contrast images were generated to examine the neural activations in response to advantageous versus disadvantageous decks: *(deckC+deckD) – (deckA+deckB)*.

For task-related data analyses, gPPI analysis was conducted to examine the functional connectivity of the MOFC during performing the IGT. The PPI method examines changes in correlation between seed regions and the rest of the brain vary as a function of the experimental manipulation, enabling analysis of task-dependent regional interactions (Kohlberg et al., 2019; McLaren et al., 2012). To generate the left and right MOFC functional networks, the seed regions were defined based on the Automated Anatomical Labeling (AAL) atlas (Tzourio-Mazoyer et al., 2002). The psychological variables were defined as the contrasts of advantageous decks, disadvantageous decks, and advantageous versus disadvantageous decks. At the single-subject level, PPI regressors were extracted from each subject. The PPI regressor consisted of the convolution of two functions, i.e., hemodynamic function convolved task regressor (advantageous versus disadvantageous decks) and the BOLD time course extracted from the seed region. This regressor was used to identify the individual effect of task-related functional connectivity. In the group-level analysis, repeated measures ANOVA was applied to examine the difference of intervention effect across the sham and tDCS groups. Familywise error (FWE) corrected cluster-size thresholds (p < 0.05, cluster size = 17 voxels) were obtained through Monte Carlo simulation (10,000 iterations), implemented in the latest version of AFNI’s 3dClustSim (Cox et al., 2017a, b).

### 2.10 Statistical analyses

Other statistical analyses were conducted in the Matlab 2014b and SPSS V22. The independent *t*-test/*Chi*-square test was used to examine the group differences in demographic, neuropsychological measurements and baseline IGT performance between sham and tDCS groups. In the task-related fMRI analysis, a repeated measures ANOVA was used to examine the interaction effect between group (sham vs. tDCS) and intervention (pre vs. post) on functional connectivity, and peak value within each significant brain region was extracted for the sham and tDCS groups. Then a partial correlation was used to examine the relationships between behavioral changes and functional connectivity changes, adjusted for gender, age, MMSE, HAMD, GM and head motion. A generalized linear model (GLM) was used to assess the relationship between changes in IGT performance (outcome) and changes in MOFC network measures (predictors), adjusting for gender, age, MMSE score, HAMD score, gray matter (GM), and head motion. The false discovery rate (FDR) was used to address multiple comparisons among model parameters.

## 3. Results

### 3.1 Baseline data analysis

This was a single-blind, randomized, sham-controlled experimental design. After baseline screening, six participants were excluded: one due to a history of long-term alcohol addiction (>10 years), one due to low educational level (< 6 years), two with HAMD scores > 16, and two due to difficulty completing the task. Eventually, forty-six participants were included in the following analyses (**Supplementary Figure 1**). There was no significant difference between the sham and tDCS groups regarding demographic data or baseline IGT (**Table 1**).

**Table 1.** Demographic, cognitive assessments and baseline IGT performance.

|  | **Sham** | **tDCS** | **p value** |
| --- | --- | --- | --- |
| Age | 64.7 ± 6.7 | 66.1 ± 8.2 | 0.55 |
| N (Male/Female) | 23 (4/19) | 23 (9/14) | 0.10 |
| Years of Education | 10.3 ± 2.7 | 11.1 ± 2.6 | 0.30 |
| MMSE | 27.0 ± 1.8 | 28.0 ± 2.2 | 0.12 |
| HAMD | 3.3 ± 4.1 | 4.1 ± 4.7 | 0.55 |
| *Baseline IGT performance* |  |  |  |
| Advantageous choice | 0.79 ± 0.18 | 0.70 ± 0.14 | 0.06 |
| Disadvantageous choice | 0.74 ± 0.16 | 0.68 ± 0.09 | 0.35 |
| Net score | 0.05 ± 0.16 | 0.02 ± 0.13 | 0.19 |
| Income | 101.7 ± 57.5 | 83.2 ± 42.7 | 0.22 |

Data are presented as means ± standard deviations. The independent t-test and Chi-square test were used to examine group differences. The one-way ANOVA was used to examine difference of task performance between sham and tDCS groups, controlled for age, gender, MMSE and HAMD. Abbreviations: HAMD, Hamilton Depression Rating Scale; IGT, Iowa Gambling Task; MMSE, Mini Mental State Examination.

### 3.2 IGT behavioral performance

To compare tDCS effect between sham and tDCS groups, a repeated measures ANOVA was applied to examine conventional behavior (advantageous choice, disadvantageous choice, and net score) adjusted for age, gender, MMSE and HAMD. In the IGT task performance, a significant interaction effect between group (sham vs. tDCS) and intervention (pre vs. post) was observed in net score (F = 4.48, p = 0.041), but not in advantageous (F = 1.36, p = 0.25) or disadvantageous choices (F = 0.24, p = 0.63) (**Figure 2A**). For cognitive training, no significant interaction effect was observed in advantageous choices (F = 0.79, p = 0.46) or disadvantageous choices (F = 0.45, p = 0.64). Although net score showed no significant interaction effect (F = 0.11, p = 0.90), the declined trend of performance may imply incrementally increased difficulty level (**Figure 2B**).

**Figure 2.**
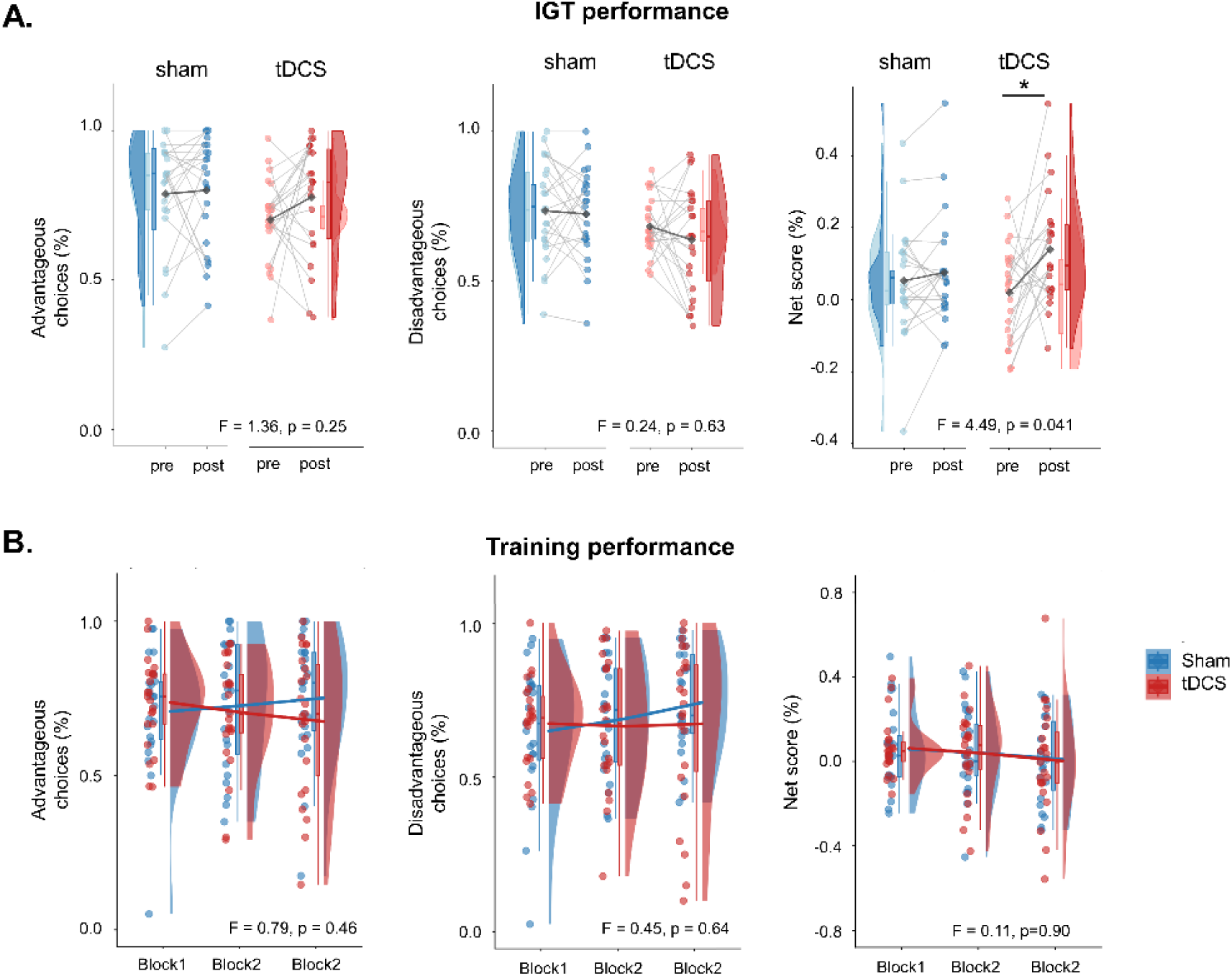
The IGT and cognitive training performance. A) Compared with the sham group, the tDCS group showed significantly increased IGT net score after stimulation. B) During the IGT training session, no significant group difference was observed across the three blocks. Paired *t*-test: *, p < 0.05.

### 3.3 IGT Computational modeling analysis

In the VPP model, remarkably interaction effects were observed in memory decay (F =160.44, p < 0.001), learning rate (F = 7896.50, p < 0.001), loss aversion (F = 998.88, p < 0.001), positive sensitivity (F = 9.71, p = 0.004), negative sensitivity (F = 9.041, p < 0.001), and preservation decay (F = 2121.25, p < 0.001), but not in consistency (F = 2.98, p = 0.092) and reinforcement learning weight (F = 3.33, p = 0.087) with FDR correction (**Figure 3**). In post hoc analyses, the tDCS intervention produced significant increases in memory decay, learning rate, loss aversion, and negative sensitivity, and a significant decrease in preservation decay. While the sham group showed significant increases in memory decay, learning rate, positive sensitivity and loss aversion, and significant decreases in negative sensitivity and perseveration decay.

**Figure 3.**
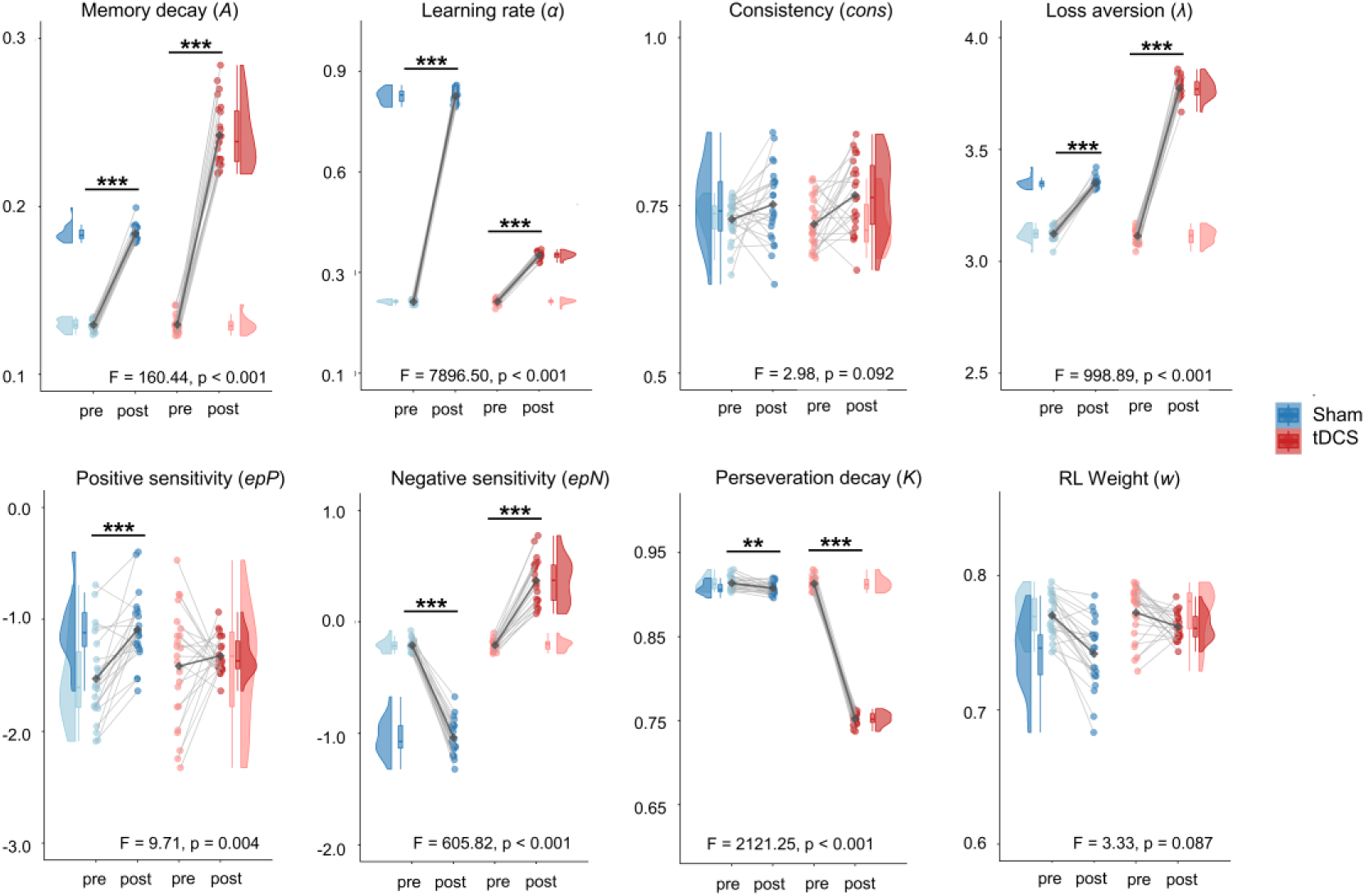
Model parameters in sham and tDCS groups. Significant interaction effects between group (sham vs. tDCS) and intervention (pre vs. post) were observed in multiple estimated parameters, including memory decay, learning rate, loss aversion, positive sensitivity, negative sensitivity and perseveration decay (FDR-corrected p < 0.05). Paired *t*-test: *, p < 0.05; **, p < 0.01; ***, p < 0.005.

### 3.4 tDCS effect on task-related MOFC network

After preprocessing neuroimaging data, the sham and tDCS participants showed no significant difference in FD (pre: t = 0.47, p = 0.64; post: t = 0.26, p = 0.80), or GM (t = 0.23, p = 0.82). In the gPPI analysis, significant interaction effects between group (sham vs. tDCS) and intervention (pre vs. post) were observed in the left and right MOFC networks, respectively (**Table 2**). In the left MOFC network, tDCS decreased connectivity strength in the right MFC (t = -2.48, p = 0.021), and increased connectivity strength in the bilateral putamen (left: t = 3.19, p = 0.004; right: t = 2.75, p = 0.012) (**Figure 4A**). In the right MOFC network, tDCS decreased connectivity strength in the right MFC (t = -2.79, p = 0.011), and increased connectivity strength in the left insula (t = 3.87, p = 0.001), right putamen (t = 3.64, p = 0.001) and left Rolandic operculum (t = 3.36, p = 0.003) (**Figure 4B**). In contrast, sham group showed significantly increased left MOFC-right MFC connectivity (t = 2.92, p = 0.008) and right MOFC-right MFC connectivity (t = 2.52, p = 0.020), and no significant changes in other connections.

**Figure 4.**
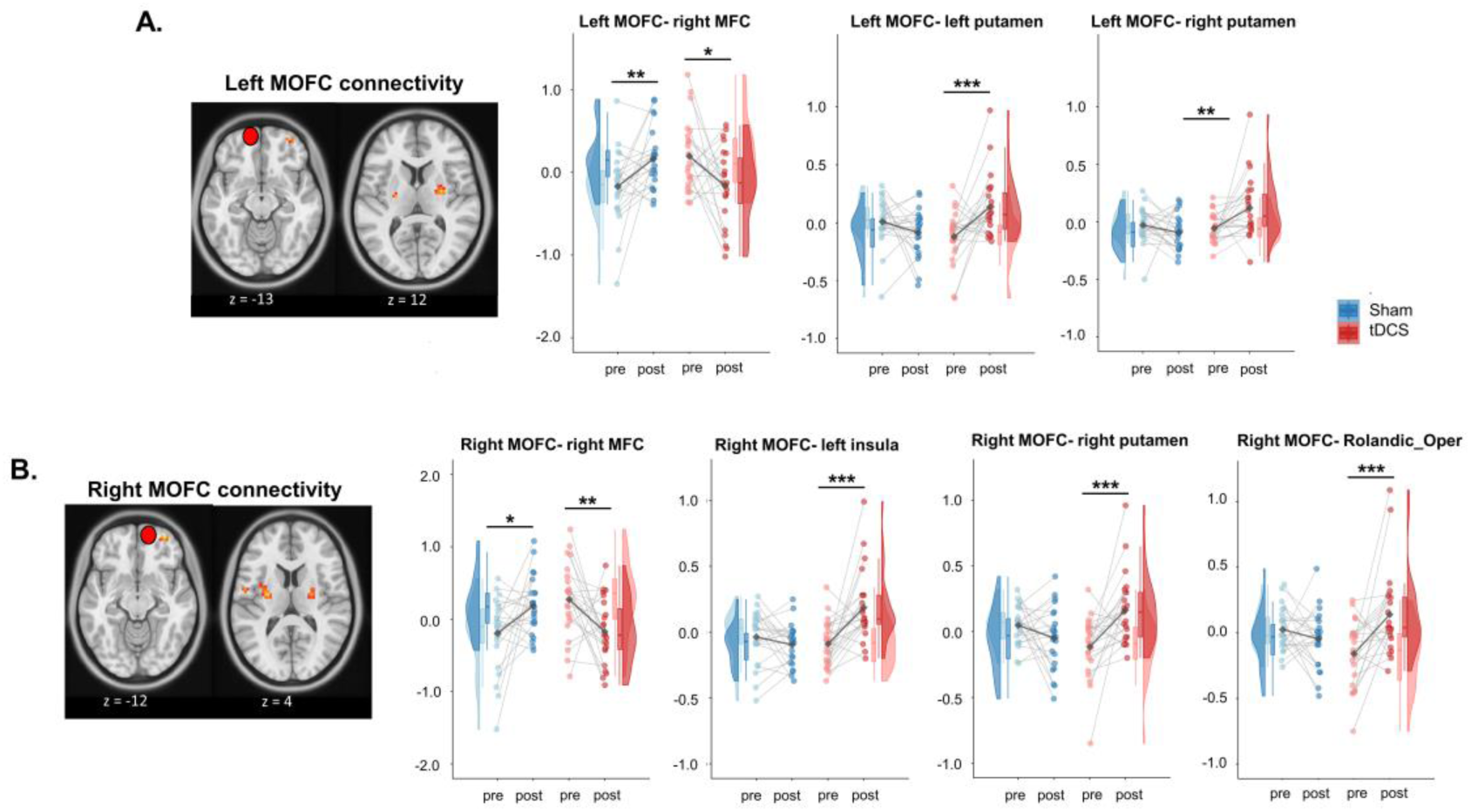
tDCS-induced group differences in MOFC connectivity. A) In the left MOFC network, significant group differences of tDCS effect were observed in the right MFC and bilateral putamen. B) In the right MOFC network, significant group differences of tDCS effect were observed in the right MFC, left insula, right putamen and left Rolandic operculum. Paired *t*-test: *, p < 0.05; **, p < 0.01; ***, p < 0.005.

**Table 2.** gPPI analysis in the MOFC network.

| Region | F value | Cluster (voxels) | MNI | BA |
| --- | --- | --- | --- | --- |
| *Left MOFC network* |  |  |  |  |
| Right MFC | 13.07 | 19 | 33, 57, -12 | 47/46/10 |
| Left putamen | 12.98 | 34 | 27, -6, 9 | N/A |
| Right putamen | 11.53 | 18 | -24, -12, 9 | N/A |
| *Right MOFC network* |  |  |  |  |
| Right MFC | 15.73 | 32 | 33, 57, -12 | 47/46/10 |
| Left insula | 15.13 | 45 | -36, 3, 9 | 48 |
| Right putamen | 14.87 | 38 | 27, -9, 9 | N/A |
| Right Rolandic operculum | 12.32 | 28 | -54, -3, 12 | 48/43 |

A Repeated measures ANOVA was performed to examine the differences of MOFC networks associated with the IGT. Abbreviations: BA, Brodmann area; MFC, middle frontal cortex; MOFC, medial orbitofrontal cortex; MNI, Montreal Neurological Institute.

### 3.5 Association between task performance and functional connectivity

In partial correlation analyses, disadvantageous choice was negatively correlated with changes in left MOFC–right putamen connectivity (r = −0.59, p = 0.036, FDR-corrected) within the tDCS group; no other significant correlations were found in either group. Within each group, a GLM was used to examine the relationship between changes in model parameters and intervention-induced connectivity changes. In the tDCS group, changes in multiple model parameters were significantly predicted by changes in the left MOFC network, but not by changes in the right mOFC network (**Table 3 and Supplementary Table 1**). In the sham group, no significant main effects of connectivity change were observed, except for right mOFC–right MFC connectivity (**Supplementary Tables 2 and 3**).

**Table 3.** GLM analysis of the left MOFC network in tDCS group.

| Model | Right MFC |  | Left putamen |  | Right putamen |  |
| --- | --- | --- | --- | --- | --- | --- |
| parameter | B | *p* | B | *p* | B | *p* |
| *A* | 0.009 | 0.14 | -0.007 | 0.63 | **0.033** | **0.018** |
| $\alpha$ | **0.007** | **0.008** | **0.013** | **0.048** | -0.011 | 0.11 |
| *cons* | -0.003 | 0.97 | -0.019 | 0.60 | 0.001 | 0.94 |
| $\lambda$ | 0.058 | 0.136 | -0.007 | 0.92 | 0.094 | 0.29 |
| *epP* | 0.11 | 0.56 | -0.69 | 0.088 | **1.04** | **0.018** |
| *epN* | 0.059 | 0.35 | -0.22 | 0.13 | **0.31** | **0.035** |
| *K* | 0.000 | 0.97 | **0.019** | **0.016** | **-0.031** | **< 0.001** |
| *w* | 0.008 | 0.18 | -0.014 | 0.35 | **0.036** | **0.018** |

Note. For each model parameter in the IGT, a GLM was applied to examine the relationship between change of parameter and change of MOFC network, adjusted for age, gender, MMSE, HAMD, gray matter and head motion. Here *p* refers to FDR-adjusted *p* value among model parameters. Abbreviations: MFC, middle frontal cortex; MOFC, medial orbitofrontal cortex.

## 4. Discussion

In the current study, we investigated the effects of mOFC-targeted tDCS combined with cognitive training on modulating risky decision-making in healthy older adults. Compared with the sham group, the tDCS group showed significantly better performance in differentiating between advantageous and disadvantageous decks. Consistently, intervention effects were observed in multiple VPP parameters, with differential changes across components. Task-related fMRI analysis revealed notable changes in the MOFC functional network, involving multiple regions including the MFC and putamen. Furthermore, tDCS-induced changes in model parameters could be predicted by changes in left MOFC connections, particularly those with the putamen. To the best of our knowledge, this is the first study examining the neural substrates of tDCS combined with cognitive training in modulating risk-taking behavior in aging.

In behavioral data analysis, significantly changed IGT net score after tDCS intervention indicated an enhanced ability of distinguishing advantageous and disadvantageous options. Consistently, a recent study has reported that OFC-tDCS could improve the IGT net score by allowing patients to attribute adequate values to risk level and choose with a better appreciation of long-term outcomes (Danon et al., 2025). León et al. reported positive improvements in risky decision-making after stimulating the right orbitofrontal cortex (León et al., 2020), while Ouellet et al. indicated enhanced decision-making abilities after stimulating either the left or right orbitofrontal cortex (Ouellet et al., 2015). Based on the “hot” and “cold” cognition model, the DLPFC is assumed to mainly be involved in “cold” cognition (e.g., executive control, working memory), while the orbitofrontal cortex is mainly involved in “hot” cognition (e.g., reward and punishment, value encoding) (Nejati et al., 2018; Salice et al., 2024; Zelazo et al., 2014). Here, since the advantageous and disadvantageous choices were not significantly changed after intervention, our finding implies that MOFC-tDCS may facilitate estimating value differences rather than value per se. Notably, there was no significant changes of net score during the IGT-based training, which may be due to the increased difficulty level across training blocks. In addition, another possibility is that tDCS may show greater effects during the offline (delayed effect) compared with online (immediate effect) (Fan et al., 2025; Friehs and Frings, 2019; Živanović et al., 2021). However, the difference between online and offline effect is beyond the scope of the present study, it would be interesting to investigate the underlying mechanism in future research.

Given the large variability of the IGT performance in older individuals, conventional task indices such as net score may not be sufficient to reveal cognitive mechanism of decision process under uncertainty. In the VPP modeling analysis, significant changes induced by tDCS were observed in multiple components. In specific, tDCS group showed significantly increased learning rate (*α*), loss aversion (*λ*), and negative sensitivity (*epN*). These alterations suggest that tDCS could accelerate learning speed and enhance loss-aversion behavior during the IGT, both of which have been found to be impaired in older adults in our previous studies (Ren et al., 2024). Interestingly, the tDCS group exhibited higher memory decay (*A*) and lower perseveration (*K*), which may have a negative effect on the decision-making process. Taken together, tDCS induced differing changes in the decision-making components, suggesting that the underlying mechanisms are much more complex than expected.

In the task-related fMRI data, the gPPI analysis showed significantly changed connectivity in both left and right MOFC networks during the IGT. Zha et. al. has reported that the OFC representation of advantageous choice was correlated to net scores in the IGT (Zha et al., 2022). Compared with the sham group, the tDCS group exhibited significantly decreased connectivity of the left MOFC-MFC network and increased connectivity of the left MOFC-putamen network. Consistently, the similar results were observed in the frontal and subcortical regions in the right MOFC network as well. Previous studies have shown that anodal tDCS effectively reduce age-related hyper-activity and hyper-connectivity in the prefrontal region (Fiori et al., 2018; Meinzer et al., 2013; Meinzer et al., 2015; Tao et al., 2021), suggesting increased neural efficiency or re-organization (i.e., less over-recruitment in response to cognitive demand). Here we speculate that the reduced MOFC-MFC coupling might represent an improved neural efficiency during cognitive task. Our previous work has demonstrated that decreased putamen activity has been found in mild cognitive impairment, leading to high risk of AD (Ren et al., 2016). An animal study has shown that disconnection between the MOFC and striatum impaired risky choice in response to varied reward probabilities through changing win-stay and lose-shift behavior (Jenni et al., 2022). Notably, our finding showed reduced disadvantageous choice associated with increased left MOFC-right putamen connectivity, highlighting the fronto-striatal interaction in risk-avoiding behavior. In aging brain, there is a general tendency to increase short-range connections and decrease long-range, reflecting the need for more local processing to compensate global inefficiency (Ferré et al., 2020; Sala-Llonch et al., 2015). Taken together, our findings suggest that MOFC-tDCS improved the brain efficiency by reducing local connectivity and enhancing global connectivity.

In tDCS intervention, the GLM analysis showed that the alterations of model components could be estimated using the left MOFC connectivity change, especially the connectivity with putamen. This result confirms the critical role of the fronto-striatal connection in regulating value-based behavior (Averbeck and O’Doherty, 2022; Zhou et al., 2023). Notably, the relationships between model parameters and functional connections were observed only in the left MOFC network (anodal stimulation) but not in the right MOFC network (cathodal stimulation). This result confirmed the effect of anodal tDCS over the left MOFC in regulating decision-making process in older adults. In line with our finding, a recent study reported that anode over the medial prefrontal cortex increased punishments-aversive behavior, whereas cathode showed no effect on participant’s behavior (Panitz et al., 2022).

Previous studies have shown consistent beneficial effects of prefrontal anodal tDCS, whereas cathodal stimulation produced smaller or no effects (Lipp et al., 2020; You et al., 2024). However, the current study only tested tDCS effect with left anodal and right cathodal stimulation over the MOFC. As previous studies have reported lateralization in PFC-related cognitive functioning, future work should consider switching electrode polarity, or place the reference electrode over alternative regions such as extracephalic positions. Therefore, our finding implies that single-session MOFC-tDCS can effectively regulate risky decision-making via the fronto-striatal connectivity.

Several limitations should be acknowledged. First, although we observed notable tDCS effects on behavioral performance and brain network, our small sample size may have limited statistical power, increasing the risk of missing potential significant findings. Therefore, future studies should validate these findings in a larger sample size. Second, the current study involved more female participants. A recent meta-analysis study systematically investigated gender differences in IGT performance, demonstrating significant differences in decision strategy and underlying brain mechanisms between male and female participants (Zanini et al., 2024). Thus, it is necessary to investigate tDCS effects on risk-taking behaviors in male and female older adults, separately. Finally, a single-session intervention may not provide maximal benefits. A consecutive stimulation design, such as 30min/day over one or two weeks, might induce greater and long-lasting tDCS effect.

## 5. Conclusions

In summary, our findings show that MOFC-tDCS combined with cognitive training significantly alters risk-taking behavior in older adults. Computational modeling analysis is more informative in characterizing the altered decision-making components. Furthermore, tDCS alters cortical and subcortical connections differently, suggesting distinct roles for these pathways in the decision-making process. Our findings provide novel insights into the use of non-invasive brain stimulation for early-stage intervention in age-related cognitive decline.

## Abbreviations

AD: Alzheimer’s disease
BART: Balloon Analogue Risk Task
DLPFC: Dorsolateral prefrontal cortex
FD: Framewise displacement
GLM: General linear model
HAMD: Hamilton Rating Scale for Depression
IGT: Iowa Gambling Task
MFC: Middle frontal cortex
MMSE: Mini-Mental State Examination
MOFC: Medial orbitofrontal cortex
tDCS: Transcranial direct current stimulation
VPP model: Values-Plus-Perseveration model

## Acknowledgements

We acknowledge Meiling Tan, Guohua Xie, and Shiwei Huang for their kind help during data collection. We appreciate all who participated in this study. This work was supported by: Natural Science Foundation of Guangdong Province 2023A1515012911 (to PR), Shenzhen Science and Technology Innovation Commission JCYJ20210324133208023 (to PR), Sanming Project of Medicine in Shenzhen SZSM201812052, National Natural Science Foundation of China 82360271, and Shenzhen Fund for Guangdong Provincial High-level Clinical Key Specialties SZGSP013.

## Data availability statement

The data generated by this study are undergoing additional analyses. The data used in this study is available from the corresponding author upon request.

## Supplementary material

Supplementary material is available online.

## References

1. Ahn, W.-Y., Vasilev, G., Lee, S.-H., Busemeyer, J.R., Kruschke, J.K., Bechara, A., Vassileva, J., 2014. Decision-making in stimulant and opiate addicts in protracted abstinence: evidence from computational modeling with pure users. Frontiers in psychology 5, 849.

2. Ahn, W.Y., Busemeyer, J.R., Wagenmakers, E.J., Stout, J.C., 2008. Comparison of decision learning models using the generalization criterion method. Cognitive science 32, 1376–1402.

3. Aksu, S., Indahlastari, A., O’Shea, A., Marsiske, M., Cohen, R., Alexander, G.E., DeKosky, S.T., Hishaw, G.A., Dai, Y., Wu, S.S., 2024. Facilitation of working memory capacity by transcranial direct current stimulation: a secondary analysis from the augmenting cognitive training in older adults (ACT) study. GeroScience 46, 4075–4110.

4. Antonenko, D., Fromm, A.E., Thams, F., Kuzmina, A., Backhaus, M., Knochenhauer, E., Li, S.-C., Grittner, U., Flöel, A., 2024. Cognitive training and brain stimulation in patients with cognitive impairment: a randomized controlled trial. Alzheimer’s research & therapy 16, 6.

5. Ashburner, J., 2007. A fast diffeomorphic image registration algorithm. Neuroimage 38, 95–113.

6. Averbeck, B., O’Doherty, J.P., 2022. Reinforcement-learning in fronto-striatal circuits. Neuropsychopharmacology 47, 147–162.

7. Busemeyer, J.R., Stout, J.C., 2002. A contribution of cognitive decision models to clinical assessment: decomposing performance on the Bechara gambling task. Psychological assessment 14, 253.

8. Cardoso, C.d.O., Carvalho, J.C.N., Cotrena, C., Bakos, D.d.G.S., Kristensen, C.H., Fonseca, R.P., 2010. Reliability study of the neuropsychological test Iowa gambling task. Journal Brasileiro de Psiquiatria 59, 279–285.

9. Carlisi, C.O., Norman, L., Murphy, C.M., Christakou, A., Chantiluke, K., Giampietro, V., Simmons, A., Brammer, M., Murphy, D.G., Mataix-Cols, D., 2017. Shared and disorder-specific neurocomputational mechanisms of decision-making in autism spectrum disorder and obsessive-compulsive disorder. Cerebral Cortex 27, 5804–5816.

10. Cauffman, E., Shulman, E.P., Steinberg, L., Claus, E., Banich, M.T., Graham, S., Woolard, J., 2010. Age differences in affective decision making as indexed by performance on the Iowa Gambling Task. Developmental psychology 46, 193.

11. Cox, R.W., Chen, G., Glen, D.R., Reynolds, R.C., Taylor, P.A., 2017a. fMRI clustering and false-positive rates. Proceedings of the National Academy of Sciences 114, E3370–E3371.

12. Cox, R.W., Chen, G., Glen, D.R., Reynolds, R.C., Taylor, P.A., 2017b. FMRI clustering in AFNI: false-positive rates redux. Brain connectivity 7, 152–171.

13. Dalley, J.W., Robbins, T.W., 2017. Fractionating impulsivity: neuropsychiatric implications. Nature Reviews Neuroscience 18, 158–171.

14. Danon, M., Perrain, R., Gorwood, P., Jollant, F., 2025. Improved decision-making in patients with mood disorders following Transcranial Direct Current Stimulation (tDCS) applied to the left orbitofrontal cortex: A proof-of-concept study. Journal of affective disorders, 119682.

15. Dedoncker, J., Brunoni, A.R., Baeken, C., Vanderhasselt, M.A., 2016. A Systematic Review and Meta-Analysis of the Effects of Transcranial Direct Current Stimulation (tDCS) Over the Dorsolateral Prefrontal Cortex in Healthy and Neuropsychiatric Samples: Influence of Stimulation Parameters. Brain Stimul 9, 501–517.

16. Di Rosa, E., Mapelli, D., Arcara, G., Amodio, P., Tamburin, S., Schiff, S., 2017. Aging and risky decision-making: New ERP evidence from the Iowa Gambling Task. Neurosci Lett 640, 93–98.

17. Fan, L., Bass, E., Klein, H., Springfield, C., Vanneste, S., Pinkham, A.E., 2025. Potential delayed positive effects of tDCS on improving introspective accuracy in social cognition in schizophrenia. Schizophrenia Bulletin, sbaf014.

18. Fein, G., McGillivray, S., Finn, P., 2007. Older adults make less advantageous decisions than younger adults: Cognitive and psychological correlates. Journal of the International Neuropsychological Society 13, 480–489.

19. Ferré, P., Jarret, J., Brambati, S.M., Bellec, P., Joanette, Y., 2020. Task-induced functional connectivity of picture naming in healthy aging: the impacts of age and task complexity. Neurobiology of language 1, 161–184.

20. Fiori, V., Kunz, L., Kuhnke, P., Marangolo, P., Hartwigsen, G., 2018. Transcranial direct current stimulation (tDCS) facilitates verb learning by altering effective connectivity in the healthy brain. NeuroImage 181, 550–559.

21. Folstein, M.F., Folstein, S.E., McHugh, P.R., 1975. “Mini-mental state”: a practical method for grading the cognitive state of patients for the clinician. Journal of psychiatric research 12, 189–198.

22. Frank, C.C., Seaman, K.L., 2023. Aging, uncertainty, and decision making—A review. Cognitive, Affective, & Behavioral Neuroscience 23, 773–787.

23. Friehs, M.A., Frings, C., 2019. Offline beats online: transcranial direct current stimulation timing influences on working memory. Neuroreport 30, 795–799.

24. Hasuzawa, S., Tomiyama, H., Murayama, K., Ohno, A., Kang, M., Mizobe, T., Kato, K., Matsuo, A., Kikuchi, K., Togao, O., 2022. Inverse association between resting-state putamen activity and Iowa gambling task performance in patients with obsessive-compulsive disorder and control subjects. Front Psychiatry 13, 836965.

25. Hultman, C., Tjernström, N., Vadlin, S., Rehn, M., Nilsson, K.W., Roman, E., Åslund, C., 2022. Exploring decision-making strategies in the Iowa gambling task and rat gambling task. Frontiers in Behavioral Neuroscience 16, 964348.

26. Indahlastari, A., Albizu, A., O’Shea, A., Forbes, M.A., Nissim, N.R., Kraft, J.N., Evangelista, N.D., Hausman, H.K., Woods, A.J., Alzheimer’s Disease Neuroimaging, I., 2020. Modeling transcranial electrical stimulation in the aging brain. Brain Stimul 13, 664–674.

27. Jenni, N.L., Rutledge, G., Floresco, S.B., 2022. Distinct medial orbitofrontal–striatal circuits support dissociable component processes of risk/reward decision-making. Journal of Neuroscience 42, 2743–2755.

28. Kohlberg, J., Preuss, A., Steiger, V., 2019. Training emotion regulation through real-time fMRI neurofeedback of amygdala activity. Neuroimage 184, 687–696.

29. León, J., Sánchez-Kuhn, A., Fernández-Martín, P., Páez-Pérez, M., Thomas, C., Datta, A., Sánchez-Santed, F., Flores, P., 2020. Transcranial direct current stimulation improves risky decision making in women but not in men: a sham-controlled study. Behavioural Brain Research 382, 112485.

30. Lipp, J., Draganova, R., Batsikadze, G., Ernst, T., Uengoer, M., Timmann, D., 2020. Prefrontal but not cerebellar tDCS attenuates renewal of extinguished conditioned eyeblink responses. Neurobiology of Learning and Memory 170, 107137.

31. Liu, Q., Cui, H., Li, J., Shen, Y., Zhang, L., Zheng, H., 2024. Modulation of dlPFC function and decision-making capacity by repetitive transcranial magnetic stimulation in methamphetamine use disorder. Translational Psychiatry 14, 280.

32. Lopez-Persem, A., Bastin, J., Petton, M., Abitbol, R., Lehongre, K., Adam, C., Navarro, V., Rheims, S., Kahane, P., Domenech, P., 2020. Four core properties of the human brain valuation system demonstrated in intracranial signals. Nature Neuroscience 23, 664–675.

33. McLaren, D.G., Ries, M.L., Xu, G., Johnson, S.C., 2012. A generalized form of context-dependent psychophysiological interactions (gPPI): a comparison to standard approaches. Neuroimage 61, 1277–1286.

34. Meinzer, M., Lindenberg, R., Antonenko, D., Flaisch, T., Flöel, A., 2013. Anodal transcranial direct current stimulation temporarily reverses age-associated cognitive decline and functional brain activity changes. Journal of Neuroscience 33, 12470–12478.

35. Meinzer, M., Lindenberg, R., Phan, M.T., Ulm, L., Volk, C., Flöel, A., 2015. Transcranial direct current stimulation in mild cognitive impairment: behavioral effects and neural mechanisms. Alzheimer’s & Dementia 11, 1032–1040.

36. Meyer, B., Mann, C., Gotz, M., Gerlicher, A., Saase, V., Yuen, K.S.L., Aedo-Jury, F., Gonzalez-Escamilla, G., Stroh, A., Kalisch, R., 2019. Increased Neural Activity in Mesostriatal Regions after Prefrontal Transcranial Direct Current Stimulation and l-DOPA Administration. J Neurosci 39, 5326–5335.

37. Mordillo-Mateos, L., Turpin-Fenoll, L., Millan-Pascual, J., Nunez-Perez, N., Panyavin, I., Gomez-Arguelles, J.M., Botia-Paniagua, E., Foffani, G., Lang, N., Oliviero, A., 2012. Effects of simultaneous bilateral tDCS of the human motor cortex. Brain Stimul 5, 214–222.

38. Nejati, V., Salehinejad, M.A., Nitsche, M.A., 2018. Interaction of the left dorsolateral prefrontal cortex (l-DLPFC) and right orbitofrontal cortex (OFC) in hot and cold executive functions: Evidence from transcranial direct current stimulation (tDCS). Neuroscience 369, 109–123.

39. Ouellet, J., McGirr, A., Van den Eynde, F., Jollant, F., Lepage, M., Berlim, M.T., 2015. Enhancing decision-making and cognitive impulse control with transcranial direct current stimulation (tDCS) applied over the orbitofrontal cortex (OFC): A randomized and sham-controlled exploratory study. Journal of psychiatric research 69, 27–34.

40. Panitz, M., Deserno, L., Kaminski, E., Villringer, A., Sehm, B., Schlagenhauf, F., 2022. Anodal tDCS over the medial prefrontal cortex enhances behavioral adaptation after punishments during reversal learning through increased updating of unchosen choice options. Cerebral Cortex Communications 3, tgac006.

41. Park, H., Kim, M., Kwak, Y.B., Cho, K.I.K., Lee, J., Moon, S.-Y., Lho, S.K., Kwon, J.S., 2022. Aberrant cortico-striatal white matter connectivity and associated subregional microstructure of the striatum in obsessive-compulsive disorder. Molecular Psychiatry 27, 3460–3467.

42. Pasion, R., Goncalves, A.R., Fernandes, C., Ferreira-Santos, F., Barbosa, F., Marques-Teixeira, J., 2017. Meta-Analytic Evidence for a Reversal Learning Effect on the Iowa Gambling Task in Older Adults. Front Psychol 8, 1785.

43. Pellicciari, M.C., Brignani, D., Miniussi, C., 2013. Excitability modulation of the motor system induced by transcranial direct current stimulation: a multimodal approach. Neuroimage 83, 569–580.

44. Power, J.D., Barnes, K.A., Snyder, A.Z., Schlaggar, B.L., Petersen, S.E., 2012. Spurious but systematic correlations in functional connectivity MRI networks arise from subject motion. Neuroimage 59, 2142–2154.

45. Ren, P., Lo, R.Y., Chapman, B.P., Mapstone, M., Porsteinsson, A., Lin, F., Initiative, A.s.D.N., 2016. Longitudinal alteration of intrinsic brain activity in the striatum in mild cognitive impairment. Journal of Alzheimer’s disease 54, 69–78.

46. Ren, P., Luo, G., Huang, J., Tan, M., Wu, D., Rong, H., 2023. Aging-related changes in reward-based decision-making depend on punishment frequency: An fMRI study. Frontiers in Aging Neuroscience 15, 1078455.

47. Ren, P., Ma, M., Zhuang, Y., Huang, J., Tan, M., Wu, D., Luo, G., 2024. Dorsal and ventral fronto-amygdala networks underlie risky decision-making in age-related cognitive decline. GeroScience 46, 447–462.

48. Sala-Llonch, R., Bartrés-Faz, D., Junqué, C., 2015. Reorganization of brain networks in aging: a review of functional connectivity studies. Frontiers in psychology 6, 663.

49. Salice, S., Antonietti, A., Colautti, L., 2024. The effect of transcranial Direct Current Stimulation on the Iowa Gambling Task: a scoping review. Frontiers in psychology 15, 1454796.

50. Samanez-Larkin, G.R., Knutson, B., 2015. Decision making in the ageing brain: changes in affective and motivational circuits. Nat Rev Neurosci 16, 278–289.

51. Summers, J.J., Kang, N., Cauraugh, J.H., 2016. Does transcranial direct current stimulation enhance cognitive and motor functions in the ageing brain? A systematic review and meta-analysis. Ageing Res Rev 25, 42–54.

52. Tang, Y., Xing, Y., Zhu, Z., He, Y., Li, F., Yang, J., Liu, Q., Li, F., Teipel, S.J., Zhao, G., 2019. The effects of 7-week cognitive training in patients with vascular cognitive impairment, no dementia (the Cog-VACCINE study): A randomized controlled trial. Alzheimer’s & Dementia 15, 605–614.

53. Tao, Y., Ficek, B., Wang, Z., Rapp, B., Tsapkini, K., 2021. Selective functional network changes following tDCS-augmented language treatment in primary progressive aphasia. Frontiers in Aging Neuroscience 13, 681043.

54. Tzourio-Mazoyer, N., Landeau, B., Papathanassiou, D., Crivello, F., Etard, O., Delcroix, N., Mazoyer, B., Joliot, M., 2002. Automated anatomical labeling of activations in SPM using a macroscopic anatomical parcellation of the MNI MRI single-subject brain. NeuroImage 15, 273–289.

55. Van Dijk, K.R., Sabuncu, M.R., Buckner, R.L., 2012. The influence of head motion on intrinsic functional connectivity MRI. Neuroimage 59, 431–438.

56. Weber, M.J., Messing, S.B., Rao, H., Detre, J.A., Thompson-Schill, S.L., 2014. Prefrontal transcranial direct current stimulation alters activation and connectivity in cortical and subcortical reward systems: A tDCS-fMRI study. Human brain mapping 35, 3673–3686.

57. Worthy, D.A., Pang, B., Byrne, K.A., 2013. Decomposing the roles of perseveration and expected value representation in models of the Iowa gambling task. Frontiers in psychology 4, 640.

58. Xu, S., Korczykowski, M., Zhu, S., Rao, H., 2013. Risk-taking and impulsive behaviors: A comparative assessment of three tasks. Social Behavior and Personality: an international journal 41, 477–486.

59. Yan, C.G., Wang, X.D., Zuo, X.N., Zang, Y.F., 2016. DPABI: Data Processing & Analysis for (Resting-State) Brain Imaging. Neuroinformatics 14, 339–351.

60. You, G., Pan, X., Li, J., Zhao, S., 2024. Effects of transcranial direct current stimulation on modulating executive functions in healthy populations: a systematic review and meta-analysis. Frontiers in Human Neuroscience 18, 1485037.

61. Zanini, L., Picano, C., Spitoni, G.F., 2024. The Iowa Gambling Task: Men and women perform differently. A meta-analysis. Neuropsychology review, 1-21.

62. Zanini, L., Picano, C., Spitoni, G.F., 2025. The Iowa Gambling Task: Men and women perform differently. A meta-analysis. Neuropsychology Review 35, 211–231.

63. Zelazo, P., Qu, L., Müller, U., 2014. Hot and cool aspects of executive function: Relations in early development. Young children’s cognitive development. Psychology Press, pp. 71–93.

64. Zha, R., Li, P., Liu, Y., Alarefi, A., Zhang, X., Li, J., 2022. The orbitofrontal cortex represents advantageous choice in the Iowa gambling task. Human brain mapping 43, 3840–3856.

65. Zhang, L., Vashisht, H., Nethra, A., Slattery, B., Ward, T., 2022. Differences in learning and persistency characterizing behavior in chronic pain for the Iowa Gambling Task: Web-based laboratory-in-the-field study. Journal of Medical Internet Research 24, e26307.

66. Zheng, X., Alsop, D.C., Schlaug, G., 2011. Effects of transcranial direct current stimulation (tDCS) on human regional cerebral blood flow. Neuroimage 58, 26–33.

67. Zhou, X., Xu, T., Zeng, Y., Zhang, R., Qi, Z., Zhao, W., Kendrick, K.M., Becker, B., 2023. The angiotensin antagonist Losartan modulates social reward motivation and punishment sensitivity via modulating midbrain-striato-frontal circuits. Journal of Neuroscience 43, 472–483.

68. Zimmerman, M., Martinez, J.H., Young, D., Chelminski, I., Dalrymple, K., 2013. Severity classification on the Hamilton depression rating scale. Journal of affective disorders 150, 384–388.

69. Živanović, M., Paunović, D., Konstantinović, U., Vulić, K., Bjekić, J., Filipović, S.R., 2021. The effects of offline and online prefrontal vs parietal transcranial direct current stimulation (tDCS) on verbal and spatial working memory. Neurobiology of Learning and Memory 179, 107398.

