## Supplementary for "Transcranial direct current stimulation combined cognitive training modulates risk-taking behavior in older adults"

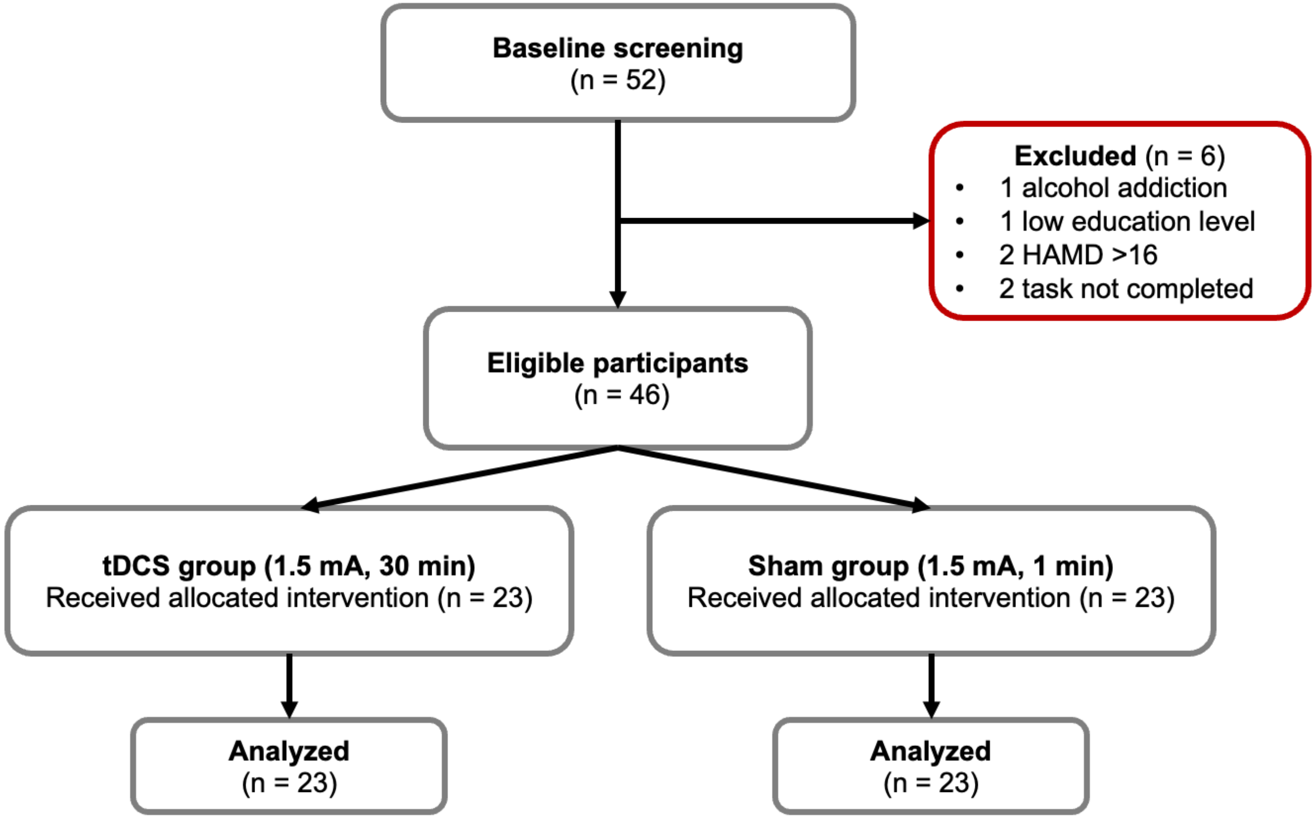


**Figure S1. Study flow diagram.**

**Table S1. GLM analysis of right MOFC network in tDCS group.**

| Model parameter | Right MFC B | Right MFC *p* | Left insula B | Left insula *p* | Right putamen B | Right putamen *p* | Right Rolandic Oper B | Right Rolandic Oper *p* |
|---|---|---|---|---|---|---|---|---|
| *A* | 0.011 | 0.080 | 0.007 | 0.70 | 0.021 | 0.48 | -0.011 | 0.58 |
| *α* | 0.005 | 0.080 | -0.011 | 0.22 | -0.007 | 0.56 | 0.013 | 0.21 |
| *cons* | -0.004 | 0.87 | -0.013 | 0.70 | -0.001 | 0.98 | -0.020 | 0.59 |
| *λ* | 0.056 | 0.080 | 0.091 | 0.41 | 0.081 | 0.56 | -0.079 | 0.58 |
| *epP* | 0.14 | 0.496 | 0.86 | 0.15 | 0.13 | 0.97 | -0.46 | 0.58 |
| *epN* | -0.007 | 0.88 | -0.204 | 0.17 | -0.204 | 0.48 | -0.014 | 0.91 |
| *K* | -0.001 | 0.88 | 0.002 | 0.15 | 0.002 | 0.97 | 0.004 | 0.66 |
| *w* | 0.009 | 0.080 | 0.032 | 0.15 | 0.011 | 0.61 | -0.12 | 0.58 |

Note. For each model parameter in the IGT, a GLM was applied to examine the relationship between change of parameter and change of MOFC network, adjusted for age, gender, MMSE, HAMD, gray matter and head motion. Here *p* refers to FDR-adjusted *p* value among model parameters. Abbreviations: MFC, middle frontal cortex; MOFC, medial orbitofrontal cortex.

**Table S2. GLM analysis of right MOFC network in sham group.**

| Model | Right MFC | | Left putamen | | Right putamen | |
|---|---|---|---|---|---|---|
| parameter | B | *p* | B | *p* | B | *p* |
| *A* | -0.007 | 0.64 | 0.000 | 0.96 | 0.003 | 0.82 |
| *α* | 0.001 | 0.95 | -0.002 | 0.96 | -0.026 | 0.70 |
| *cons* | -0.032 | 0.64 | 0.079 | 0.92 | -0.117 | 0.56 |
| *λ* | -0.035 | 0.65 | 0.044 | 0.96 | -0.095 | 0.72 |
| *epP* | -0.33 | 0.64 | -0.755 | 0.51 | 1.328 | 0.82 |
| *epN* | 0.101 | 0.64 | -0.101 | 0.96 | 0.075 | 0.28 |
| *K* | -0.003 | 0.65 | -0.002 | 0.96 | -0.002 | 0.82 |
| *w* | 0.007 | 0.65 | -0.019 | 0.96 | 0.031 | 0.70 |

Note. For each model parameter in the IGT, a GLM was applied to examine the relationship between change of parameter and change of MOFC network, adjusted for age, gender, MMSE, HAMD, gray matter and head motion. Here *p* refers to FDR-adjusted *p* value among model parameters. Abbreviations: MFC, middle frontal cortex; MOFC, medial orbitofrontal cortex.

**Table S3. GLM analysis of right MOFC network in sham group.**

| Model | Right MFC | | Left insula | | Right putamen | | Right Rolandic Oper | |
|---|---|---|---|---|---|---|---|---|
| parameter | B | *p* | B | *p* | B | *p* | B | *p* |
| *A* | -0.001 | 0.73 | -0.007 | 0.72 | 0.002 | 0.76 | 0.010 | 0.83 |
| *α* | -0.013 | 0.32 | -0.030 | 0.72 | -0.019 | 0.59 | 0.026 | 0.83 |
| *cons* | -0.024 | 0.50 | -0.001 | 0.99 | 0.043 | 0.59 | 0.049 | 0.88 |
| *λ* | 0.012 | 0.73 | 0.099 | 0.72 | 0.021 | 0.76 | 0.005 | 0.97 |
| *epP* | -0.065 | 0.73 | 0.67 | 0.72 | -0.218 | 0.62 | -0.617 | 0.83 |
| *epN* | 0.097 | 0.35 | -0.097 | 0.87 | -0.132 | 0.59 | -0.009 | 0.97 |
| *K* | -0.003 | 0.50 | -0.003 | 0.87 | 0.005 | 0.59 | -0.004 | 0.88 |
| *w* | **0.019** | **0.024** | 0.019 | 0.72 | -0.018 | 0.59 | -0.010 | 0.88 |

Note. For each model parameter in the IGT, a GLM was applied to examine the relationship between change of parameter and change of MOFC network, adjusted for age, gender, MMSE, HAMD, gray matter and head motion. Here *p* refers to FDR-adjusted *p* value among model parameters. Abbreviations: MFC, middle frontal cortex; MOFC, medial orbitofrontal cortex.
